# Superagers resist typical age-related white matter structural changes

**DOI:** 10.1101/2024.01.25.577227

**Authors:** Marta Garo-Pascual, Linda Zhang, Meritxell Valentí-Soler, Bryan A. Strange

**Author notes:** **Corresponding authors:** (M. Garo-Pascual) and (B.A. Strange). **Conflict of interest statement:** The authors declare no competing financial interest.

## Abstract

Superagers are elderly individuals with the memory ability of people 30 years younger and provide evidence that age-related cognitive decline is not inevitable. In a sample of 64 superagers (mean age 81.9; 59% women) and 55 typical older adults (mean age 82.4; 64% women) from the Vallecas Project, we studied, cross-sectionally and longitudinally over 5 years with yearly follow-ups, the global cerebral white matter status as well as region-specific white matter microstructure assessment derived from diffusivity measures. Superagers and typical older adults showed no difference in global white matter health (total white matter volume, Fazekas score, and lesions volume) cross-sectionally or longitudinally. However, analyses of diffusion parameters revealed better white matter microstructure in superagers than in typical older adults. Cross-sectional differences showed higher fractional anisotropy (FA) in superagers mostly in frontal fibres and lower mean diffusivity (MD) in most white matter tracts, expressed as an anteroposterior gradient with greater group differences in anterior tracts. FA decrease over time is slower in superagers than in typical older adults in all white matter tracts assessed, which is mirrored by MD increases over time being slower in superagers than in typical older adults in all white matter tracts except for the corticospinal tract, the uncinate fasciculus and the forceps minor. The better preservation of white matter microstructure in superagers relative to typical older adults supports resistance to age-related brain structural changes as a mechanism underpinning the remarkable memory capacity of superagers, while their regional ageing pattern is in line with the last-in-first-out hypothesis.

**SIGNIFICANCE STATEMENT:** Episodic memory is one of the cognitive abilities most vulnerable to ageing. Although memory normally declines with age, some older people may have memory performance similar to that of people 30 years younger, and this phenomenon is often conceptualised as superageing. Understanding the superager phenotype can provide insights into mechanisms of protection against age-related memory loss and dementia. We studied the white matter structure of a large sample of 64 superagers over the age of 80 and 55 age-matched typical older adults during 5 years with yearly follow-ups showing evidence of slower age-related changes in the brains of superagers especially in protracted maturation tracts, indicating resistance to age-related changes and a regional ageing pattern in line with the last-in-first-out hypothesis.

## INTRODUCTION

Ageing is a dynamic process involving functional and structural brain changes. One of the cognitive functions most vulnerable to ageing is episodic memory, the ability to retrieve our personal experiences (Glisky, 2007). Pathological deterioration of episodic memory is a feature of Alzheimer’s disease, the leading cause of dementia. Yet episodic memory can also be robust to age-related changes and this phenomenon has been conceptualised and studied under the definition of “superagers”. Superagers are older adults with the episodic memory of a healthy adult 20-30 years younger (Cook et al., 2017; Gefen et al., 2015; Harrison et al., 2018; Harrison et al., 2012; Sun et al., 2016). Structural and functional neuroanatomical characterisation of superagers may reveal the neural substrates of successful episodic memory ageing and, thus, provide insight into how it is possible to age without episodic memory impairment. In this study we focused on structural parameters of white matter health to extend our previous work on the grey matter signature of a group of superagers form the Vallecas Project cohort (Garo-Pascual et al., 2023).

White matter undergoes changes with ageing, white matter volume decreases, microstructural properties are lost, and lesions accumulate (Cox et al., 2016; Davis et al., 2009; de Leeuw et al., 2001; Westlye et al., 2010; Ylikoski et al., 1995). These changes are regionally heterogeneous, being greater in anterior than posterior brain regions (Davis et al., 2009; Kochunov et al., 2007; O’Sullivan et al., 2001; Pfefferbaum et al., 2005; Sullivan and Pfefferbaum, 2006). This occurs in conjunction with changes of white matter microstructural properties in the thalamic radiations and association fasciculi (Cox et al., 2016; Slater et al., 2019). This white matter ageing pattern inverts the sequence of myelination early in life and supports the last-in-first-out hypothesis (Raz, 2000) since white matter tracts that first experience the effects of ageing, like the thalamic radiations and association fibres, also show protracted maturation in early life.

White matter loss with ageing is associated with worsening cognitive performance affecting processing speed, primarily impairing executive functions (Kennedy and Raz, 2009; Tubi et al., 2020). Episodic memory function in the cognitively healthy elderly is also negatively associated with white matter microstructural properties of the uncinate, inferior and superior longitudinal fasciculus, thalamic radiations, and dorsal cingulum bundle (Lockhart et al., 2012; Sasson et al., 2013; Ziegler et al., 2010). White matter microstructure has already been studied in cohorts of successful episodic memory agers, specifically in cohorts between 60-80yo, showing better properties in superagers in the corpus callosum and the right superior longitudinal fasciculus (Kim et al., 2020).

We studied the brain white matter status and white matter microstructure proxies in a sample of 64 superagers and 55 typical older adults that are over 80yo to characterise brain white matter in an older age range of superagers that is, to our knowledge, currently unexplored. We approached this study with a cross-sectional and longitudinal characterisation of 1) global white matter status and 2) white matter microstructure derived from diffusion tensor imaging parameters. In our previous study, which characterised grey matter volumes of the same sample of superagers compared to typical older adults (Garo-Pascual et al., 2023), we concluded that superagers express a resistance to age-related brain changes as manifested in greater grey matter volume and slower atrophy in the medial temporal lobe and motor thalamus compared to typical older adults. In the current study, we hypothesised that the superager brain would show resistance to age-related white matter changes and would have better global white matter status and preserved white matter microstructure –higher fractional anisotropy (FA) and lower mean diffusivity (MD)– in anterior tracts especially the anterior thalamic radiation and association fibres in comparison to typical older adults as these are the more vulnerable tracts to age-related changes.

## MATERIALS AND METHODS

### Participants

The sample of superagers and typical older adults used in this study were selected from the single-centre community based Vallecas Project, an ongoing longitudinal cohort established in Madrid (Spain). The 1,213 participants of the Vallecas Project were all of Caucasian ethnicity, community-dwelling individuals between 70 to 85 years-old, independent in activities of daily living with a survival expectancy of at least 4 years and without any neurological or psychiatric disorders (Olazarán et al., 2015). All participants provided written informed consent, and the project was approved by the Ethics Committee of the Instituto de Salud Carlos III. We applied criteria for superagers and typical older adults to the Vallecas Project cohort based on the definition of a superager as a person aged 80 years or older with the episodic memory of a person 20-30 years younger. The selection criteria for this analysis focused on five aspects including age, episodic memory performance, cognitive performance in non-memory domains, availability of MRI scans, and stability of episodic memory. Both candidates for the superager and the typical older adult group were 79.5 years or older when their episodic memory was screened with the free delayed recall score on the verbal memory free and cued selective reminding test. For participants to be considered as superagers they were required to perform at or above the mean of the score of adults aged 50-56 years with the same education attainment and typical older adults were required to score within one standard deviation from the mean of the normative values for their age and education attainment in the Spanish NEURONORMA project (Peña-Casanova et al., 2009). Complete details on the selection of superagers and typical older adults from the Vallecas project are described elsewhere (Garo-Pascual et al., 2023).

### MRI data acquisition

MRI images were acquired using a 3 Tesla MRI (Sigma HDxt GEHC, Waukesha, USA) with a phased array 8 channel head coil. T1-weighted images (3D fast spoiled gradient echo with inversion recovery preparation) were collected using a TR of 10ms, TE of 4.5ms, FOV of 240mm and a matrix size of 288×288 with slice thickness of 1mm, yielding a voxel size of 0.5 x 0.5 x 1 mm. Diffusion-weighted images were single-shot SE-EPI, with TR 9200ms, TE 80ms, b-value 800s/mm2 and 21 gradient directions, FOV 240mm and matrix size 128 x 128 with slice thickness of 3mm. T2-FLAIR (image attenuated inversion recovery) images were acquired with TR 9000 ms, TE 130 ms, TI 2100 ms, FOV 24 mm, slice thickness 3.4 mm.

### Brain white matter volume and white matter lesions volume

Brain white matter volume and white matter lesions volume were extracted from the segmentation of T1-weighted images using CAT12.7 toolbox (https://neuro-jena.github.io/cat) implemented in SPM12 (version r6225; https://www.fil.ion.ucl.ac.uk/spm) (Ashburner and Friston, 2005). This pipeline was run for cross-sectional and longitudinal analyses, with the latter including scans from visit 1 to visit 6. Total intracranial volume (TIV) was also extracted using this protocol for analytical purposes as a covariate. White matter lesions are typically detected as hyperintense radiological observations in T2-FLAIR images. Here, however, we computed the volume of white matter lesions from T1-weighted images using the CAT12 toolbox, which provides a similar performance compared to existing methods of white matter hyperintensity detention from T2-FLAIR data (Dahnke et al., 2019).

### Fazekas score

The Fazekas scale (Fazekas et al., 1987) quantifies brain white matter hyperintensities from MRI data with a scale as 0 = absence, 1 = focal lesions, 2 = start of confluent lesions and 3 = diffuse affectation in a region ± U-shaped fibres. For our cohort, the lesions were graded by a radiologist blinded to the subject’s group using T2-FLAIR images.

### White matter tract-based spatial statistics (TBSS) of diffusivity measures

For preprocessing of diffusion-weighted images, FSL was used (http://fsl.fmrib.ox.ac.uk/fsl/fslwiki) and the pipeline included a motion and eddy current correction, the extraction of non-brain voxels and ends with the calculation of voxel-wise diffusion maps —FA and MD— for each participant. Both FA and MD are derived from the eigenvalues of the diffusion tensor captured by diffusion-weighted images; FA measures the directionality of water diffusion, while MD averages the diffusivity of water molecules in the three directions of the space reflecting tissue constraints. Individual diffusion maps were then used in the TBSS pipeline using the FMRIB toolbox (http://fsl.fmrib.ox.ac.uk/fsl/fslwiki) (Smith et al., 2006). The general outline of the process is 1) FA individual maps were non-linearly registered to standard space (FMRIB58_FA template) (Andersson et al., 2007); 2) a mean FA image was created by averaging all co-registered FA maps; and 3) individually aligned images were projected onto the mean FA skeleton, representing the centres of all tracts common to the study sample (visual inspection was required to set a threshold of mean FA at 0.25 to include non-skeleton voxels) and skeletonised images were used for voxel-wise analysis. Diffusivity maps for MD were generated by applying the same steps detailed above. For cross-sectional analysis, diffusivity maps for FA and MD were entered into separate general linear models (GLM) to compare differences between the superager and the control group. TIV, age, gender and years of education were used as covariates. We conducted whole-brain analyses using a Threshold Free Cluster Enhancement (TFCE) approach with 5000 permutations (default parameters E = 0.5 and H = 2). Significant results are reported at a FWE-corrected level of *P* < 0.05. To visualise our results we used the multimodal analysis and visualisation tool (MMVT) (Felsenstein et al., 2019). The same preprocessing pipeline and GLM was built for additional diffusivity measures including mode of anisotropy, axial and radial diffusivity (Figure 1-2). While MD averages the diffusivity of water molecules in the three directions of the space, axial diffusivity reflects the diffusion of water molecules in the parallel orientation to the axonal bundle and radial diffusivity averages the two perpendicular diffusivity directions. Mode of anisotropy is mathematically orthogonal to FA and reflects the geometrical properties of the directionally of water diffusion (i.e., linear, or planar directionality). FA and MD values were also explored longitudinally replicating with longitudinal scans the same preprocessing steps described above and further used for a regions of interest (ROI)-based analysis conducted by averaging the FA and MD values from 18 ROIs described in the JHU-ICBM thr25 atlas (Hua et al., 2008; Wakana et al., 2007) (Figure 3-1, Figure 3-2). The statistical model is specified in the statistical analysis section.

**Figure 1.**
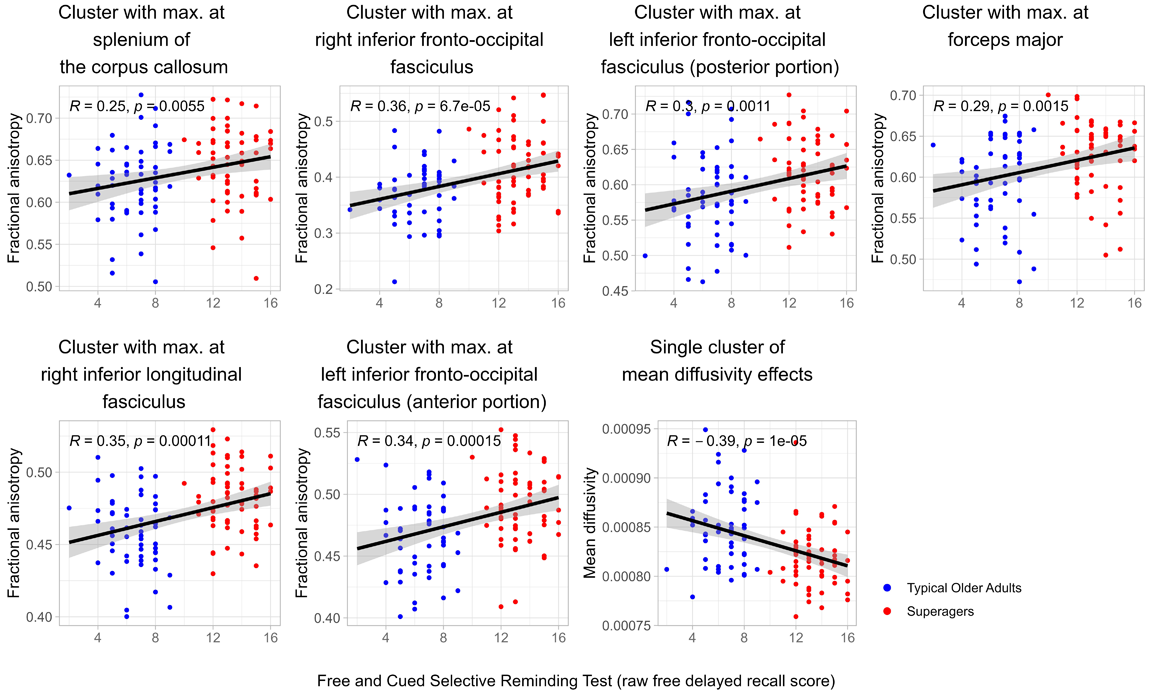
Better white matter microstructure in superagers, particularly in frontal white matter tracts, compared to typical older adults. A. Superagers show higher fractional anisotropy than typical older adults in bilateral frontal tracts and the anterior thalamic radiation (warm colours, *P* < 0.05 FWE-corrected). B. Lower mean diffusivity (MD) is found in superagers compared to typical older adults in an extensive network (cold colours, *P* < 0.05 FWE-corrected). C. Significantly greater MD group differences in the anterior half of the brain as indicated by the linear fit (blue line) of the parameter estimates (β) of the contrast MD higher in typical older adults than in superagers as a function of anteroposterior axis coordinates (positive Montreal Neurologic Initiative (MNI) coordinates for the anterior portion of the brain). See Extended Data Figure 1-1 for the correlation between FA and MD vales and episodic memory performance and Figure 1-2 for group differences in additional diffusivity measures. Mean β and ± standard error are plotted. A, anterior; L, left; R, right; P, posterior; FWE-corr p, family-wise corrected *p*-value.

**Figure 2.**
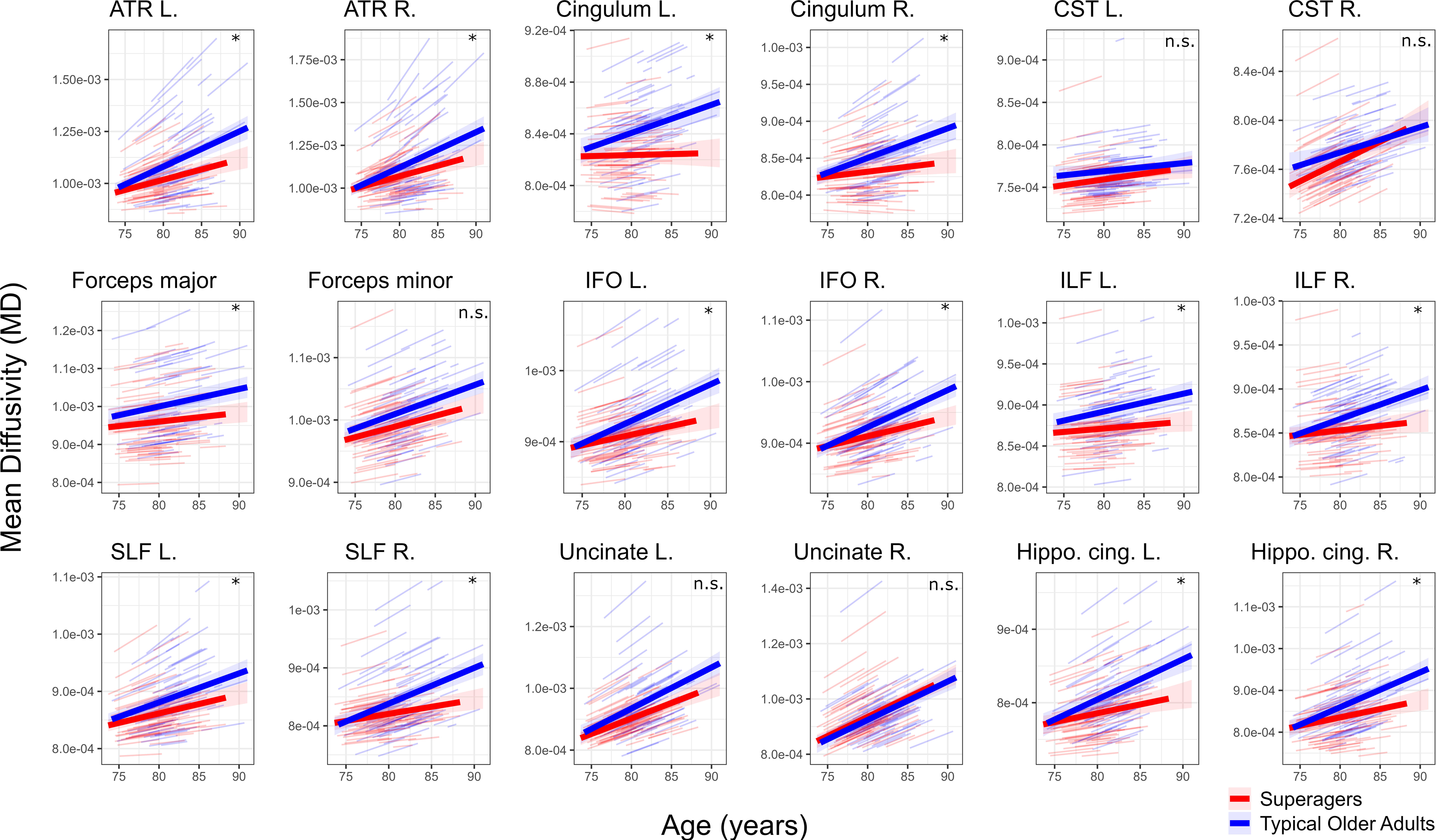
Longitudinal evolution of white matter volume and white matter lesions volume in superagers and typical older adults. **A.** Predicted trajectories of total brain white matter volume over time were plotted for superagers (red line) and typical older adults (blue line) with respective shaded areas indicating the 95% confidence interval and individual trajectories in black, showing no difference at baseline or atrophy rate between groups. **B.** Accumulation over time of white matter lesions measured as white matter lesion volume. There was no baseline difference between groups and longitudinal trajectories between groups were no longer significantly different after exclusion of outliers, three typical older adults and a superager indicated in grey. White matter volumes and white matter lesion volumes have been adjusted by total intracranial volume in the statistical model and for illustration purposes. Age was scaled in the statistical model, but raw values are shown for illustration purposes. See Extended Data Figure 2-1 for further details of the statistical models.

**Figure 3.**
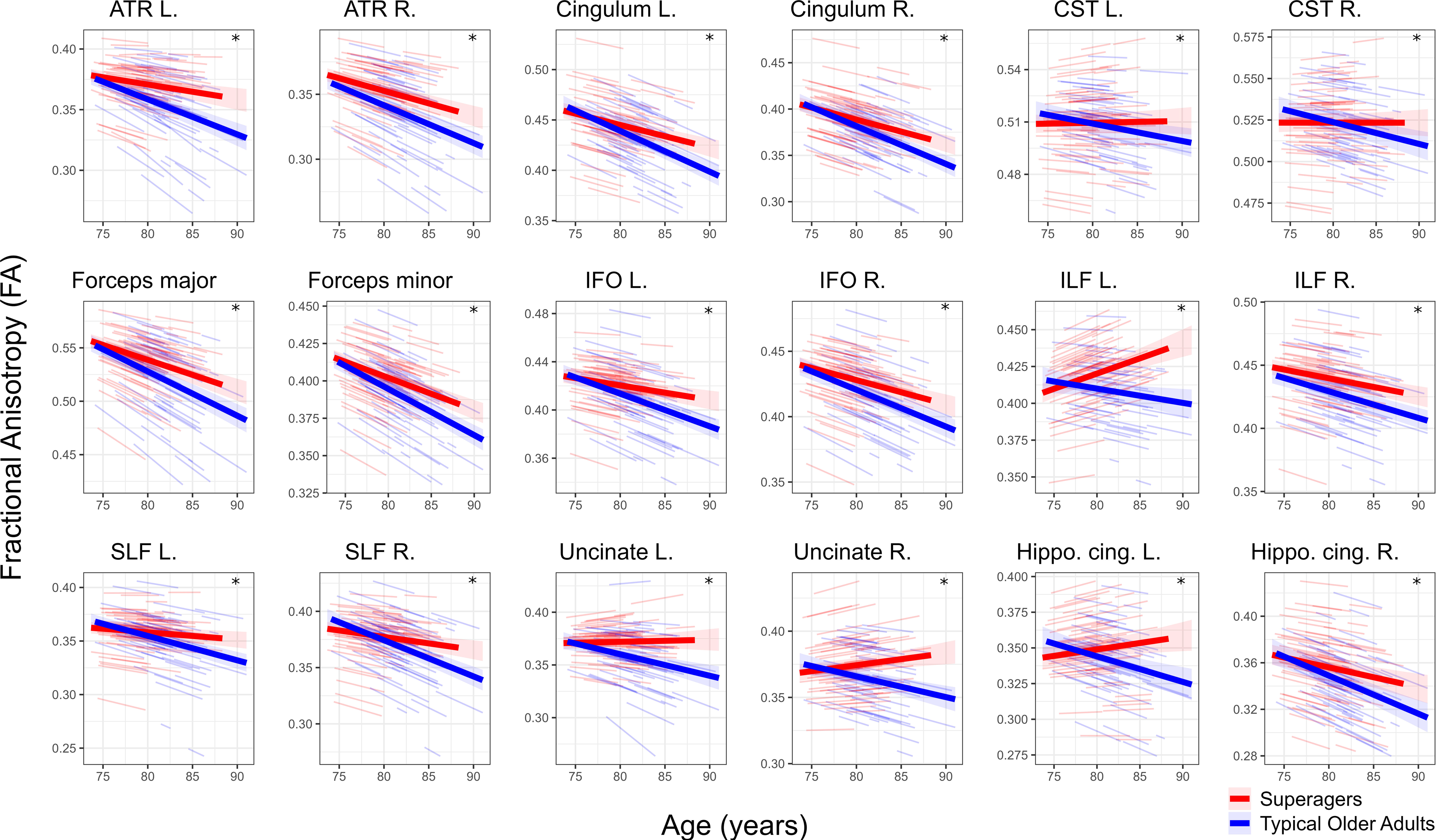
Longitudinal changes in diffusivity measures between superagers and typical older adults. **A.** Longitudinal differences in fractional anisotropy. Superagers show a slower decrease of fractional anisotropy compared to typical older adults in an extended network of white matter tracts (shaded regions) (*P* < 0.05 FWE-corrected). See Extended Data Figures 3-1 and 3-2 for ROI-based analyses. **B.** Longitudinal differences in mean diffusivity. Superagers show a slower increase of mean diffusivity compared to typical older adults in an extended network of white matter tracts (shaded regions) (*P* < 0.05 FWE-corrected). See Extended Data Figures 3-3 and 3-4 for ROI-based analyses. **C.** JHU-ICBM atlas labels of white matter tracts were used to map the significant effects shown in the rest of the panels. Note that the significant effects shown in A. and B. are not constrained to white matter since the fractional anisotropy and mean diffusivity maps were not limited to white matter skeleton. FWE-corr p, family-wise error *p*-value.

### Longitudinal diffusivity analysis in SPM

Whole-brain voxel-wise analyses testing longitudinal group differences in two measures derived from diffusion-weighted imaging sequences —FA and MD— were carried out using SPM12 (version r6225; https://www.fil.ion.ucl.ac.uk/spm) and FSL (https://fsl.fmrib.ox.ac.uk/fsl/fslwiki/) (Jenkinson et al., 2012). The preprocessing of diffusion-weighted images was conducted in FSL as described in the previous section. We performed eddy current correction, brain segmentation to exclude non-brain voxels and calculation of FA and MD parameters with the FMRIB toolbox (https://fsl.fmrib.ox.ac.uk/fsl/fslwiki). The resulted FA and MD maps were normalised to standardise Montreal Neurological Institute (MNI) space using the TBSS pipeline (Smith et al., 2006), a non-linear registration to set individual’s maps into the standard template FMRB58_FA. *Randomise*, the FSL function that builds GLM, does not support reliable longitudinal analysis, so the preprocessed data was further analysed in SPM similarly to previous authors (Lei et al., 2012).

The normalised FA and MD maps generated in FSL were then smoothed in SPM12 using a 6 mm FWHM Gaussian kernel. In the longitudinal toolbox in CAT12, separate GLM models were specified for FA and MD. Age at each MRI acquisition was included as a covariate interacting with the group factor. A masking threshold of 0.1 was applied to FA images to remove effects out of the brain. No masking threshold was used in MD images since the MD values have a low order of magnitude. These voxel-wise analyses were conducted using TFCE approach with 5000 permutations and default parameters (E = 0.5 and H = 2) using the TFCE tool (version r223) from CAT12 toolbox in SPM12 (https://www.neuro.uni-jena.de/tfce). Significant results are reported at FWE-corrected level of *P* < 0.05. The neuroanatomical loci were reported according to the Mori and the JHU-ICBM thr25 atlas (Hua et al., 2008; Wakana et al., 2007) and Mango software (http://rii.uthscsa.edu/mango/) was used to produce the figure.

### Statistical analysis

Cross-sectional group comparisons for white matter volume and white matter lesions volume were conducted with an analysis of covariance with TIV as covariate. Categorical data was evaluated with a Chi-squared test or Fisher exact test. Differences in the longitudinal trajectories of neuropsychological variables, white matter volume, white matter lesions volume and Fazekas scores (computed as numeric due to the accumulative nature of the scale) and ROI-based FA and MD values were studied with a linear mixed effects model built with the lme4 package in R (Bates et al., 2015). In these linear mixed effects models, white matter volume and white matter lesions volume were adjusted by TIV; scaled age, group and the interaction between scaled age and group were the fixed factors; and the random intercept and the random slope were included. We excluded from the longitudinal analysis of white matter lesions volume four outliers (three typical older adults and a superager) informed by the Bonferroni outlier test of the car package in R (Fox and Weisberg, 2019) that considers the longitudinal trajectory of the linear mixed effects model. Group differences in the anteroposterior gradient of MD were explored in the cross-sectional analysis. The brain map of the unthresholded parameter estimates (β) of the comparison MD in typical older adults minus MD in superagers was sliced in the coronal plane every 5mm in the anteroposterior axis creating 31 slices. We then estimated the average β in each slice and fit a linear regression to this value as a function of the anteroposterior axis Montreal Neurologic Initiative (MNI) coordinates. Whole brain significant results are reported at a TFCE corrected threshold as described above. Significant results are reported at a false discovery rate (FDR) corrected level of *P* < 0.05. All statistical analysis described were performed in R 4.0.2 (https://www.r-project.org/).

## RESULTS

A sample of 64 superagers and 55 typical older adults were identified in the Vallecas Project cohort with no significant differences in age or sex (Table 1). Superagers outperformed typical older adults in the neuropsychological selection criteria variables (Table 1), however their longitudinal evaluation showed no significant group by time interaction in the free delay recall score of the free and cued selective reminding test (t(1, 597) = 1.60, *P* = 0.11), the digit symbol substitution test (t(1, 478) = 0.86, *P* = 0.40) and the 15-item Boston naming test (t(1, 482) = 1.32, *P* = 0.19), whereas the significant group by time interaction in the animal fluency test (t(1, 600) = 2.13, *P* = 0.03) indicates a slower decline in superagers compared to typical older adults (Table 1-1).

**Table 1.**
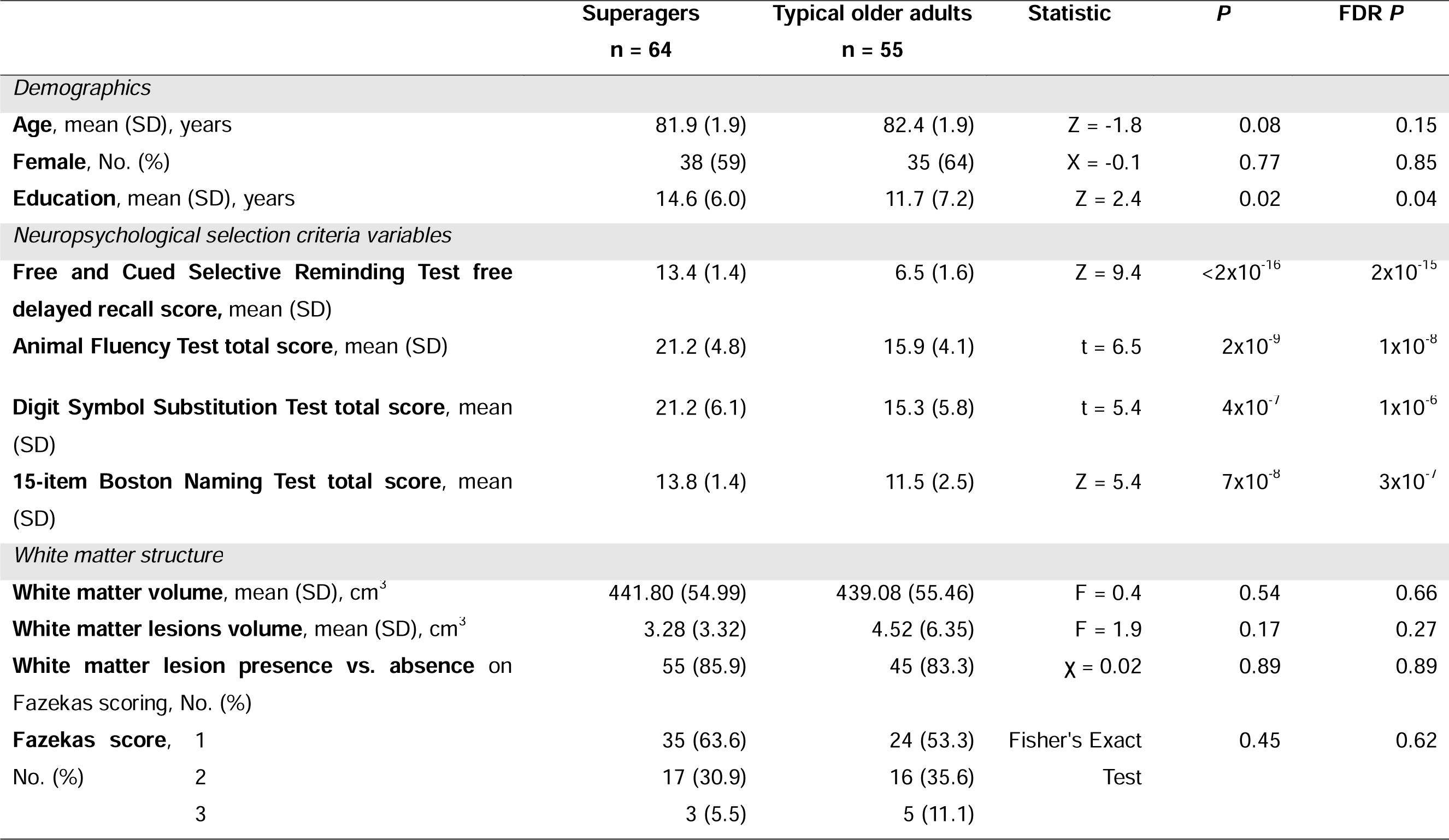
Demographic and cross-sectional white matter volume and white matter lesion differences between superagers and typical older adults. Volumetric group differences were calculated with total intracranial volume included as a covariate. The mean and standard deviation (SD) reported in the table correspond to raw values. Group differences in the Fazekas scale were assessed with a Chi-square test and Fisher’s exact test. See Extended Data Table 1-1 for the longitudinal evolution of the neuropsychological variables. FDR *P*, False Discovery Rate *p*-value; *P*, *p*-value.

White matter status of superagers and typical older adults was compared cross-sectionally and longitudinally over five years using three parameters to assess the general status of white matter: 1) total brain white matter volume, 2) volume of white matter lesions extracted automatically from T1-weighted images and 3) the Fazekas score (Fazekas et al., 1987), a radiological scale for quantifying the amount of white matter T2 hyperintense lesions (see Methods). This general white matter status assessment was complemented with a regionally specific approach to test for voxel-wise group cross-sectional and longitudinal differences in two diffusivity measures, FA and MD.

### Cross-sectional white matter structural differences between superagers and typical older adults

Superagers and typical older adults showed no cross-sectional differences in total white matter volume (F(1, 115) = 0.4, *P* = 0.54) (Table 1) or in the volume of white matter lesions (F(1, 115) = 2.0, *P* = 0.17) (Table 1). The Fazekas scores revealed that a similar proportion of superagers (85.9%) and typical older adults (83.3%) have white matter T2 hyperintense lesions (χ = 0.02, *P* = 0.89) (Table 1). This high prevalence of white matter lesions is in accordance with observations from other elderly cohorts (American Psychiatric Association, 1994; de Leeuw et al., 2001). There were no between-group differences in the degree of these lesions (*P* = 0.45) (Table 1). We observed a significant correlation between white mater lesion volume and the Fazekas score (Pearson’s r = 0.73, *P* < 0.0001).

We next adopted a regionally specific approach to test for cross-sectional voxel-wise group differences in FA and MD. Better white matter microstructure in terms of diffusivity translates into higher FA and lower MD values. We observed higher FA in superagers than typical older adults mainly in fontal regions of the inferior fronto-occipital fasciculus, anterior thalamic radiation, right inferior longitudinal fasciculus, right forceps minor and left forceps major (*P* < 0.05 FWE-corrected) (Figure 1A). Lower MD values were found in superagers compared to typical older adults in an extensive network comprising the forceps major and minor, superior and inferior longitudinal fasciculus, inferior fronto-occipital fasciculus, anterior thalamic radiation, and cingulum bundle (*P* < 0.05 FWE-corrected) (Figure 1B). The anteroposterior gradient of these group effects (higher MD in typical older adults than superagers) was tested by fitting a linear regression model of the parameter estimate (β) of this contrast as a function of anteroposterior axis coordinates. We observed a significant effect (β (t(29) = 5.31, *P* < 0.0001)) supporting stronger MD group differences in the anterior portion of the brain (Figure 1B). The average FA and MD values of the significant clusters in the voxel-wise group contrast were correlated with the episodic memory performance in the free and cued selective reminding test (Figure 1-1). Additional diffusivity measures including axial and radial diffusivity and mode of anisotropy were explored (Figure 1-2) and support the above results of superior white matter microstructural properties in the superager brain.

### Longitudinal white matter structural differences between superagers and typical older adults

Longitudinal assessment of white matter structure, both for the general status metrics and for the regional approach on FA and MD, was performed over 5 years with yearly follow-ups (median number of follow-up visits was 5.0 (IQR 5.0-6.0) for superagers and 5.0 (4.5-6.0) for typical older adults). The longitudinal evolution of total white matter volume suggest similar atrophy rates in superagers and typical older adults as the group by time interaction is not significant (t(1, 578) = 0.2, *P* = 0.81) (Figure 2, Figure 2-1). The longitudinal load of white matter lesions volume over time is significantly slower in superagers compared to typical older adults (t(1, 578) = 2.4, *P* = 0.02) but this group by time interaction did not survive the exclusion of outliers (three typical older adults and a superager) (t(1,558) = 1.6, *P* = 0.11, β(SE) superager: 0.69 (0.14), β(SE) typical older adult: 1.01 (0.15)) (Figure 2, Figure 2-1). The longitudinal evolution of lesions degree in the Fazekas scale revealed no between-group differences (t(1, 578) = 0.3, *P* = 0.80) (Figure 2-1). Thus, of the three global parameters assessed, no major differences in white mater status were found between superagers and typical older adults cross-sectionally or longitudinally.

Region-specific diffusivity measures were studied longitudinally over five years with yearly follow-up scans using a voxel-wise approach and complementary ROI-based analyses (extended data). We observed that FA decreases significantly slower in superagers compared to typical older adults in all white matter tracts described in the JHU-ICBM atlas (Figure 3), and diffuse but significant effects were found in the cingulum, hippocampal cingulum, and uncinate fasciculus bilaterally (*P* < 0.05 FWE-corrected) (Figure 3A). ROI-based analyses yielded a significantly slower FA decrease in superagers compared to typical older adults in all white matter tracts assessed (Figure 3-1, Figure 3-2). The increase of MD over time was significantly slower in superagers compared to typical older adults in all tracts from the JHU-ICBM atlas with similar differences bilaterally and diffuse but significant effects in the cingulum (*P* < 0.05 FWE-corrected) (Figure 3B). ROI-based analyses revealed different group longitudinal trajectories in all white matter tracts except for the corticospinal tract, the uncinate fasciculus and forceps minor (Figure 3-3, Figure 3-4). Altogether, these results indicate a resistance to age-related changes in white matter microstructure in superagers compared to typical older adults by showing a slower decrease of FA and a slower increase in MD over time.

## DISCUSSION

Assessment of global cerebral white matter status indicated that superagers and typical older adults have similar white matter health cross-sectionally and longitudinally since no group differences were found in total brain white matter volume, white matter lesions volume and the Fazekas score. However, differences in diffusivity measures consistent with better white matter microstructure in superagers than typical older adults were found cross-sectionally and longitudinally. Cross-sectional differences show higher FA in superagers mostly in frontal fibres and lower MD in most white matter tracts following an anteroposterior gradient with greater group differences in anterior regions, both FA and MD values correlate with episodic memory performance in the whole sample. The decrease in FA over time is slower in superagers than typical older adults in all white matter tracts assessed and the increase of MD overtime is slower in superagers than typical older adults in all white matter tracts except for the corticospinal tract, the uncinate fasciculus and the forceps minor.

The regional study of diffusivity measures—FA and MD— confirms, firstly, better white matter microstructural properties in superagers than in typical older adults, both cross-sectionally and longitudinally, and, secondly, outlines regional brain patterns associated with ageing. Cross-sectionally, the greatest differences between groups for both FA and MD accumulate in the anterior part of the brain in line with the existing evidence that the anterior portion of the brain is more vulnerable to the effects of ageing (Davis et al., 2009; Kochunov et al., 2007; O’Sullivan et al., 2001; Pfefferbaum et al., 2005; Sullivan and Pfefferbaum, 2006). Although greater group differences in MD were found in the anterior areas, they were not constrained to the anterior portion like most FA effects. MD is a more sensitive parameter to age-related changes than FA (Cox et al., 2016), and this could explain its larger group differences. These marked group white matter differences in the anterior part of the brain, rather than the temporal, contrast with grey matter volume differences observed in medial temporal areas (Garo-Pascual et al., 2023). This result might suggest that the prefrontal cortex of superagers exerts more efficient top-down control over medial temporal regions mediating more successful episodic memory function (Dobbins et al., 2002; Simons and Spiers, 2003; Szczepanski and Knight, 2014) by, for example, improving the retrieval of appropriate memories an supressing the inappropriate ones (Anderson et al., 2016; Tomita et al., 1999).

Longitudinally, extensive group differences in white matter microstructure were found in most white matter tracts. However, ROI-based analysis revealed that MD trajectories over time were similar for both groups in the corticospinal tract, the uncinate fasciculus and the forceps minor. The absence of longitudinal differences between groups in the corticospinal tract and the forceps minor is of particular interest, as these are some of the most robust white matter tracts to the effects of ageing (Cox et al., 2016; Slater et al., 2019), supporting the last-in-first-out hypothesis (Raz, 2000). Therefore, the ageing trajectories between superagers and typical older adults mainly differ in association fibres and the anterior thalamic radiation — which are the most vulnerable to age-related changes (Cox et al., 2016; Slater et al., 2019)— reinforcing the idea that superagers exhibit a resistance mechanism to age-related changes (Garo-Pascual et al., 2023) and suggesting that the differential white matter status between groups has not been established in an early developmental stage. Indeed, longitudinal ROI-based FA and MD trajectories suggest equivalent values in both groups at around age 75, consistent with previous findings in grey matter volume (Garo-Pascual et al., 2023), at a time when superagers already outperformed typical older adults in episodic memory function. This suggests that the cognitive profile of superagers is established before reaching the criterion age and before structural brain differences are evident. The super-ageing phenotype may be dictated by a resistance versus a resilience mechanism, opposing concepts (Arenaza-Urquijo and Vemuri, 2018) that in the context of healthy ageing reflect as avoidance of age-related changes versus coping with age-related changes respectively (Garo-Pascual et al., 2023). Therefore, in brain structural terms, resistance to age-related changes translates into the better preservation of brain structure in superagers than in typical older adults consistent with our white matter microstructural findings. Further evidence that resistance is the most plausible mechanism for superagers is its comparison with a group of middle-aged adults which, in our case, is not within the studied population demographic of the Vallecas Project cohort.

Fazekas scores revealed a high proportion of participants, whether superagers or typical older adults, with hyperintense white matter T2 lesions (∼85%). Likewise, both groups experienced a longitudinal accumulation of white matter lesions, indexed by both the Fazekas scale and white matter lesion volume load, measures that were correlated in our sample. This high prevalence of white matter lesions is in line with observations from other elderly cohorts (American Psychiatric Association, 1994; de Leeuw et al., 2001), as is the increasing load of white matter lesions over time (Ylikoski et al., 1995). The correlation between the Fazekas scale and the volume of white matter lesions in our sample is also consistent with individuals in other elderly cohorts (Cedres et al., 2020; Valdes Hernandez Mdel et al., 2013; van Straaten et al., 2006). The absence of group differences in the prevalence and cumulative progression of white matter brain lesions reveals that these features are not only present in healthy ageing individuals but also occur in superageing. Superagers may be then showing resilience to white matter lesions in concurrence with resistance to age-related structural changes (including white matter microstructure and grey matter volume (Garo-Pascual et al., 2023)) as the primary protective ageing mechanism for maintenance of memory function.

The similar global white mater health between groups based on volumetric and radiological metrics contrasts with the better white matter microstructure of superagers relative to typical older adults observed on the regional study of diffusivity measures. This apparent discrepancy may have two explanations that are not mutually exclusive, the higher sensitivity of regional-based approaches over global measures and the differential ageing pattern of white matter volume and diffusivity measures. The white matter volume lifespan pattern exhibits an inverted U-shape peaking during the 5th-6th decade (Walhovd et al., 2011; Westlye et al., 2010), while diffusivity measures —including FA and MD— follow the same parabolic pattern but peak around two decades earlier (Westlye et al., 2010). The time window in which we assessed our population is closer to the peak of white matter volume maturation than to the peak of diffusivity measures. Therefore, the shorter time between white matter volume maturation and our assessment could explain the similar group ageing trajectories despite finding a divergent ageing pattern in diffusivity measures.

White matter lesions (Haller et al., 2013) and age-related changes in white matter diffusion properties (Song et al., 2003; Song et al., 2005) underlie axonal and/or myelin degeneration yielding negative consequences for cognitive function (de Groot et al., 2000; Prins and Scheltens, 2015). The age-related accumulation of white matter lesions affects processing speed, mainly impairing executive function and, to a lesser extent, the memory domain (Prins and Scheltens, 2015; Tubi et al., 2020). Changes in white matter microstructure accounted by diffusivity measures have also a deleterious effect on memory performance (Goldstein et al., 2009). Poor white matter health has been associated with a vascular aetiology, as the prevalence of cardiovascular disease is a risk factor for the enlargement of white matter lesions (Debette and Markus, 2010; Launer et al., 2000) and the accumulation of vascular risk factors is associated with diffusivity parameters of impaired white matter microstructure (Ingo et al., 2021). Superagers showed lower prevalence of hypertension and glucose disorders than typical older adults (Garo-Pascual et al., 2023). However, they do not show differences in other cardiovascular risk factors (de Bruijn and Ikram, 2014) like high cholesterol, smoking status, obesity, diet —quantified as weekly frequency of food groups and adherence to Mediterranean diet— and physical activity (Garo-Pascual et al., 2023). Thus, the better white matter health in the brains of superagers relative to typical older adults could be explained by a higher burden of vascular risk factors in typical older adults, although not all cardiovascular risk factors support this speculation.

In summary, the better overall preservation of white matter microstructure in the brain of superagers supports resistance to age-related changes as their most plausible protective mechanism for maintenance of memory function, in line with our previous results from structural analyses of grey matter of the superaging brain (Garo-Pascual et al., 2023). The regional ageing pattern identified a better preservation of white matter microstructural properties in superagers at the anterior portion of the brain and in those tracts with a protracted maturation which, according to the last-in-first-out hypothesis, are more vulnerable to age-related changes (Raz, 2000). The similar properties between superagers and healthy older adults in early developing white matter tracts may indicate that the superageing phenotype is not established during early development but is rather the result of a different ageing process in which vascular health might play an influential role.

## Author contributions

MGP, LZ and BAS contributed to the conceptualisation of the study, MVS contributed with data curation, MGP and, LZ, contributed to the methodology and the investigation work of the study, BAS was responsible for the supervision of this work, MGP drafted the original manuscript and MGP, LZ, and BAS reviewed and edited the manuscript.

## Data sharing

The Vallecas Project data collection is expected to be completed by the end of 2023. Anonymised data can be accessed upon request at.

## Acknowledgments

We thank the participants of the Vallecas Project and the staff of the CIEN Foundation. This work was supported by the CIEN Foundation and the Queen Sofia Foundation, as well as by a grant from the Spanish Ministry of Science and Innovation (PID2020-119302RB-I00) to BAS. MGP was supported by a MAPFRE-Queen Sofia Foundation scholarship. LZ was supported by a grant from the Alzheimer’s Association (2016-NIRG-397128) to BAS.

## EXTENDED DATA

**Table 1-1. Longitudinal trajectories of the neuropsychological tests used for selection criteria.** Linear fits of performance trajectories of **A.** the free and cued selective reminding test raw free delayed recall score, **B.** the animal fluency test total score, **C.** the digit symbol substitution test total score and **D.** the 15-item Boston naming test total score over time are plotted for superagers (red solid line) and typical older adults (blue solid line). Respective shaded areas indicate the 95% confidence interval and individual participant trajectories are also plotted (thin lines). The threshold for episodic memory performance in superagers (at or above the mean score of a 53-year-old person with the same education level) is indicated with a dashed line in the free and cued selective reminding test plot. For the rest of the tests, typical older adults did not have a set criterion while superagers had to perform within one standard deviation from the mean for their age and education. In linear mixed effects models assessing group differences in longitudinal neuropsychological performance, age was scaled but raw values are shown for illustration purposes. The significant interaction between scaled age and group (*P* < 0.05) is indicated with an asterisk (*), otherwise as non-significant (n.s.).

**Figure 1-1. Association between fractional anisotropy and mean diffusivity values and episodic memory performance in superagers and typical older adults.** Fractional anisotropy and mean diffusivity values were extracted from the cross-sectional diffusivity images averaging the values of the significant clusters from group contrasts, six clusters from the contrast higher fractional anisotropy in superagers vs. typical older adults and a single large cluster form the contrast higher mean diffusivity in typical older adults vs. superagers. The correlation with the raw free delayed recall score of the free and cued reminding test was performed with a Peason’s correlation test and the correlation coefficient (R) and the *p*-value (*p*) of each test are shown in the plot. The clusters are named under the white matter tract of the JHU-ICBM atlas where their global maximum is located.

**Figure 1-2.** Radial and axial diffusivity and mode of anisotropy cross-sectional differences between superagers and typical older adults. A. **Lower radial diffusivity and B.** lower axial diffusivity is found in superagers compared to typical older adults in an extensive network (cold colours) comprising all the tracts described in JHU-ICBM atlas consisting with the group effects in mean diffusivity. **C.** Higher mode of anisotropy is found in superagers than typical older adults in a small part of the left inferior longitudinal fasciculus and forceps major (warm colours). This result shows that, despite the group differences in fractional anisotropy that reflect a stronger directionality of water diffusivity in the frontal tracts of superager’s brains, there is no major group differences in the shape of this directionality. (*P* < 0.05 FWE-corrected). A, anterior; FWE-corr p, family-wise error *p*-value; L, left; R, right and P, posterior.

**Figure 2-1. Longitudinal evolution of white matter volume, white matter lesions volume and Fazekas score.** Coefficients (β) correspond to the three linear mixed effects model predicting the longitudinal evolution of total brain white matter volume, brain white matter lesion volume and Fazekas score progression respectively. In the three independent models, age, group and the interaction between age and group were fixed effects (scaled age was introduced in the model) and the random intercept and slope were also considered in the model. In the white matter lesions volume analysis four outliers were excluded (three typical older adults and a superager). White matter volume and white matter lesions volume were adjusted by total intracranial volume (TIV). *P*, *p*-value; SD, standard deviation; SE, standard error.

**Figure 3-1. ROI-based longitudinal trajectories of white matter fractional anisotropy (FA).** Longitudinal group differences were studied in 18 white matter tracts or regions of interest (ROIs) from the JHU-ICBM atlas. Individual trajectories are plotted together with the group average predicted trajectory. Shaded areas indicate 95% confidence interval. A linear mixed effects model was used to predict average FA in each of the ROIs with group, scaled age and the interaction between the two as fixed factors, the random intercept and slope were included in the model. Age was scaled in the statistical model, but raw values are shown for illustration purposes. All ROIs show a significant (*P* < 0.05) interaction between age and group (indicated with an asterisk (*)). ATR, anterior thalamic radiation; CST, corticospinal tract; Hippo. cing., hippocampal cingulum; IFO, inferior fronto-occipital fasciculus; ILF, inferior longitudinal fasciculus; L, left; R, right; SLF, superior longitudinal fasciculus.

**Figure 3-2. ROI-based longitudinal analysis of white matter fractional anisotropy (FA).** Longitudinal group differences were studied in 18 white matter tracts or regions of interest (ROIs) from the JHU-ICBM atlas with a linear mixed effects model to predict average FA in each of the ROIs with group, scaled age and the interaction between the two as fixed factors, the random intercept and slope were included in the model. ATR, anterior thalamic radiation; β, coefficients; Corr. *P*, false-discovery-rate corrected *p*-value; CST, corticospinal tract; Hippo. cing., hippocampal cingulum; IFO, inferior fronto-occipital fasciculus; ILF, inferior longitudinal fasciculus; L, left; *P*, *p*-value; R, right; SE, standard error; SLF, superior longitudinal fasciculus.

**Figure 3-3. ROI-based longitudinal trajectories of white matter mean diffusivity (MD).** Longitudinal group differences were studied bilaterally in 18 white matter tracts or regions of interest (ROIs) from the JHU-ICBM atlas. Individual trajectories are plotted together with the group average predicted trajectory. Shaded areas indicate 95% confidence interval. A linear mixed effects model was used to predict average MD in each of the ROIs with group, scaled age and the interaction between the two as fixed factors, the random intercept and slope were included in the model. Age was scaled in the statistical model, but raw values are shown for illustration purposes. ROIs where the interaction between age and group is significant (*P* < 0.05) are indicated with an asterisk (*), otherwise as non-significant (n.s.). ATR, anterior thalamic radiation; CST, corticospinal tract; Hippo. cing., hippocampal cingulum; IFO, inferior fronto-occipital fasciculus; ILF, inferior longitudinal fasciculus; L, left; R, right; SLF, superior longitudinal fasciculus.

**Figure 3-4. ROI-based longitudinal analysis of white matter mean diffusivity (MD).** Longitudinal group differences were studied in 18 white matter tracts or regions of interest (ROIs) from the JHU-ICBM atlas with a linear mixed effects model to predict average MD in each of the ROIs with group, scaled age and the interaction between the two as fixed factors, the random intercept and slope were included in the model. ATR, anterior thalamic radiation; β, coefficients; Corr. *P*, false-discovery-rate corrected *p*-value; CST, corticospinal tract; Hippo. cing., hippocampal cingulum; IFO, inferior fronto-occipital fasciculus; ILF, inferior longitudinal fasciculus; L, left; *P*, *p*-value; R, right; SE, standard error; SLF, superior longitudinal fasciculus.

---

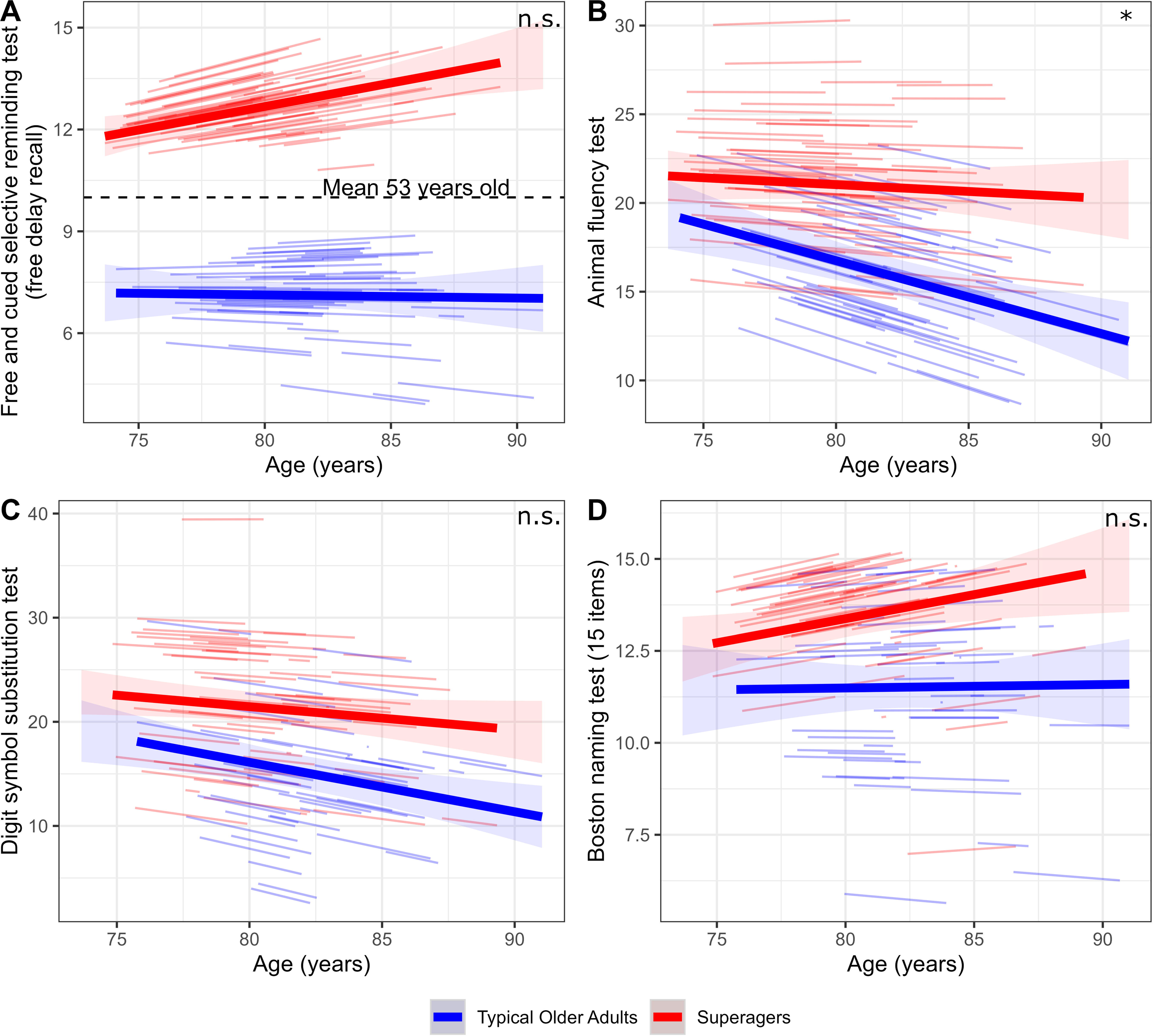

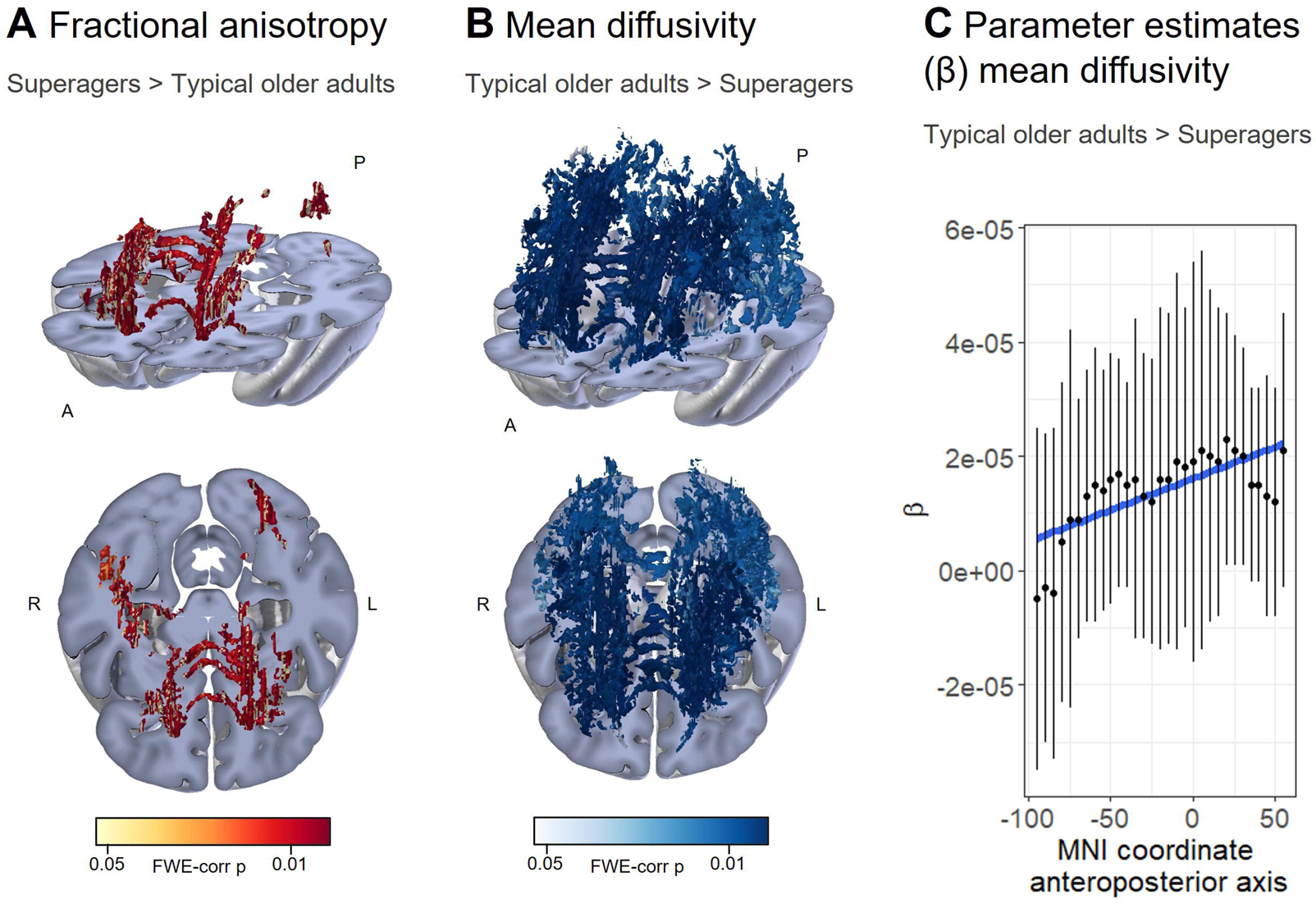

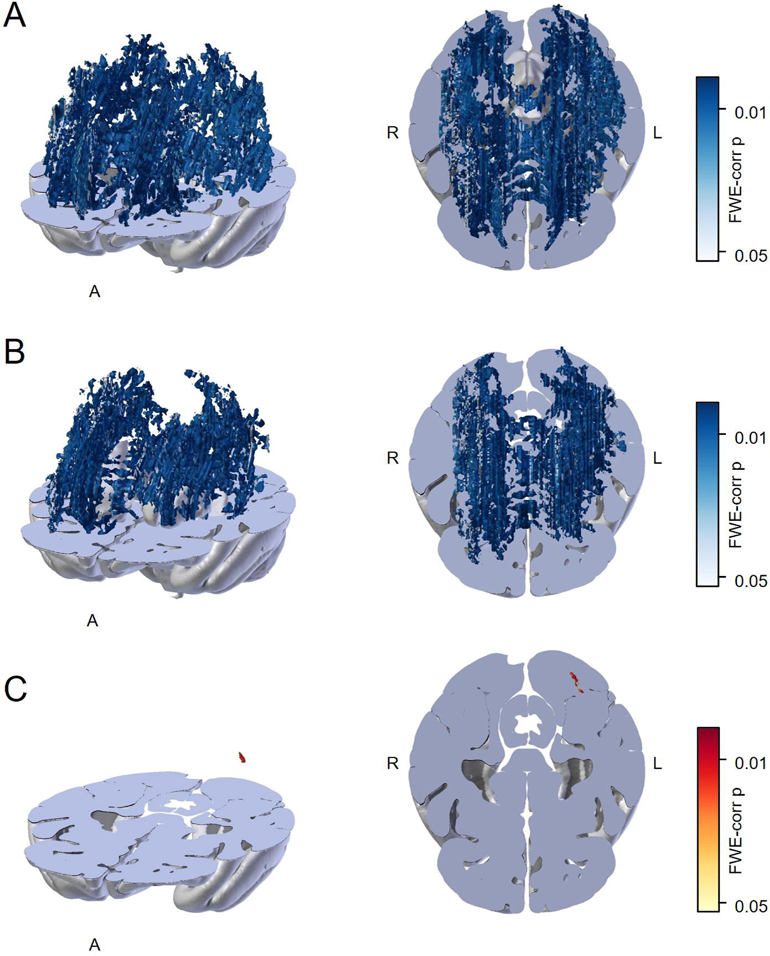

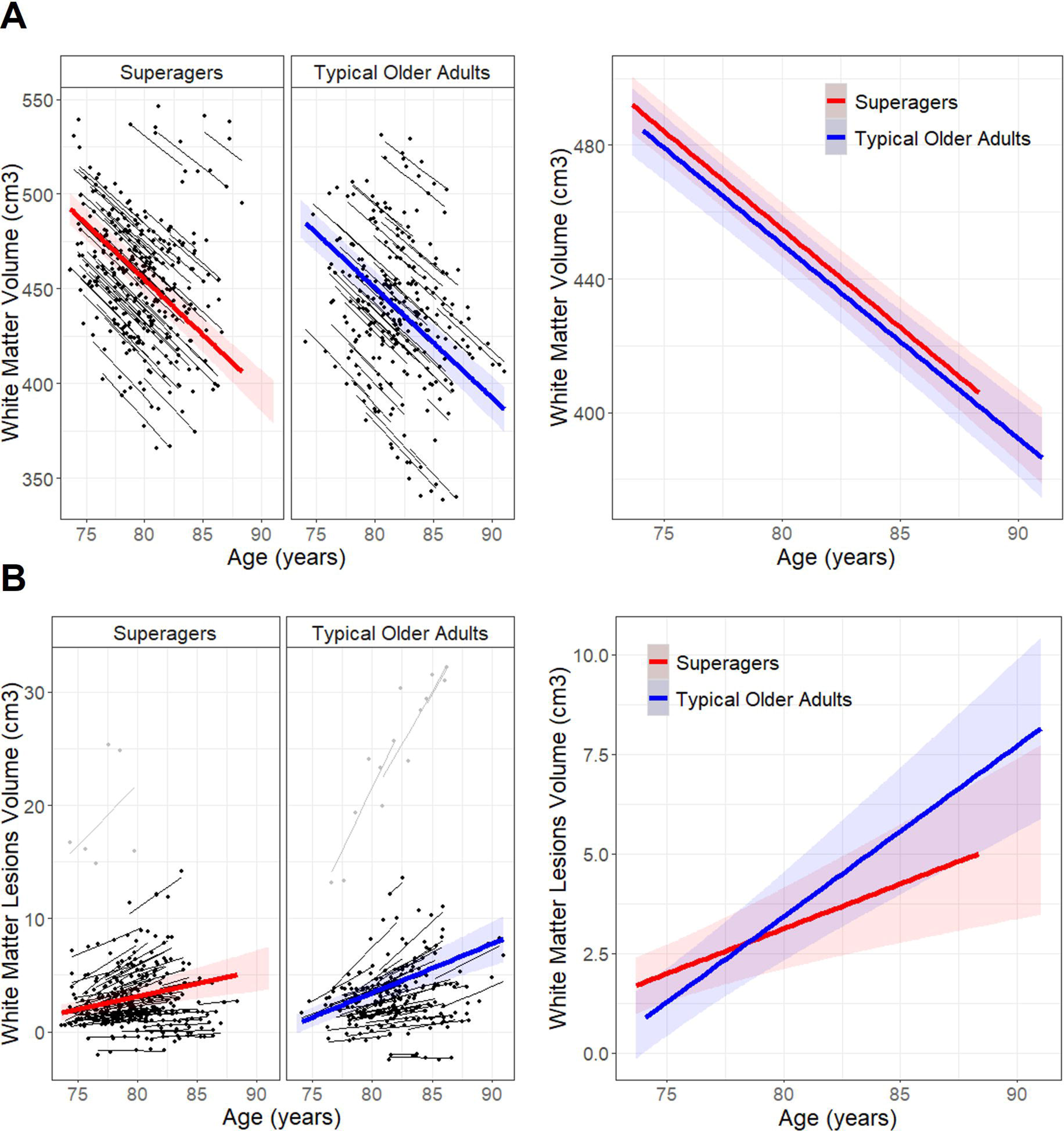

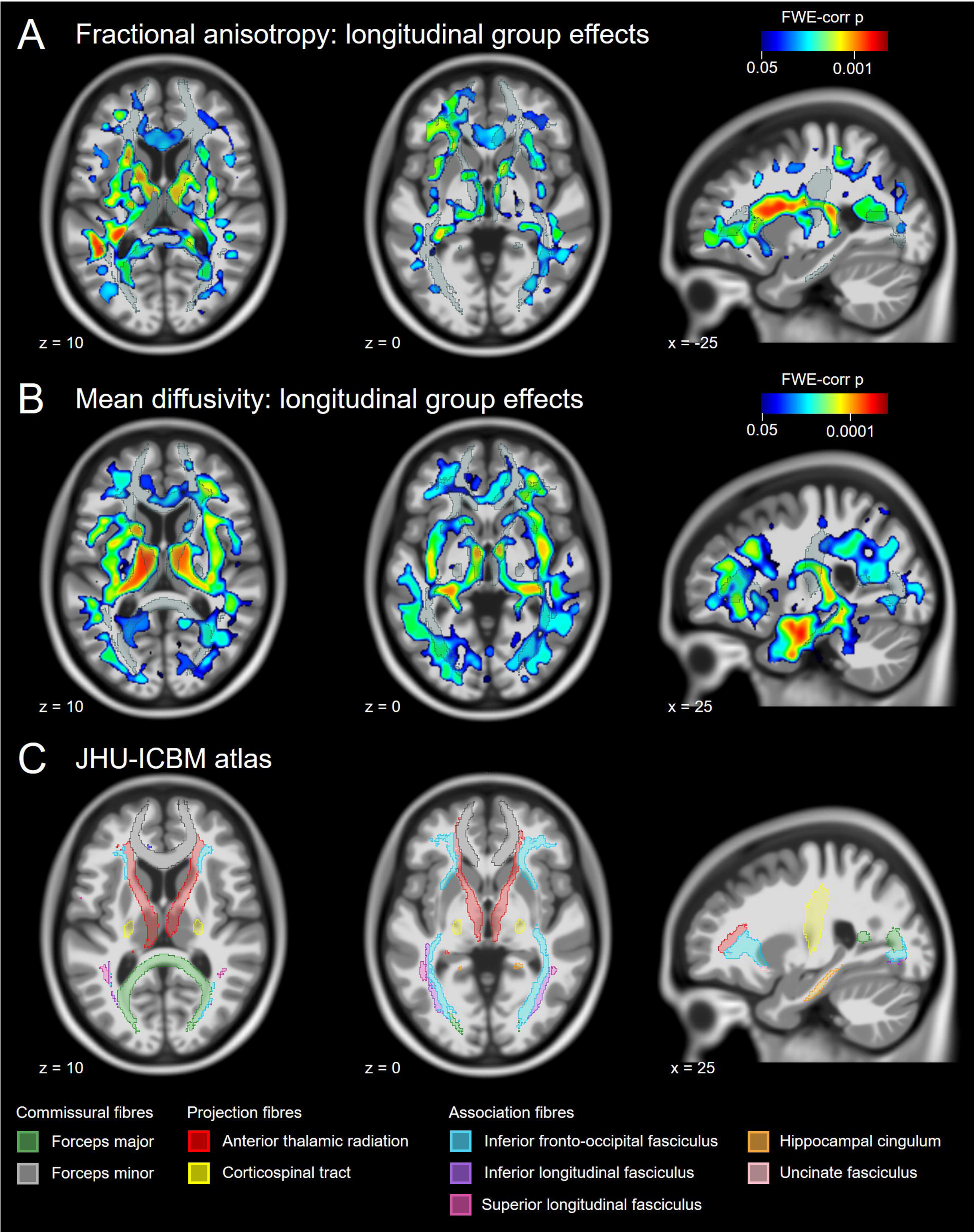

## Notes

### Competing Interest Statement

The authors have declared no competing interest.

