## Extended data legends for "Superagers resist typical age-related white matter structural changes"

**Figure 2-1. Longitudinal evolution of white matter volume, white matter lesions volume and Fazekas score.** Coefficients (β) correspond to the three linear mixed effects model predicting the longitudinal evolution of total brain white matter volume, brain white matter lesion volume and Fazekas score progression respectively. In the three independent models, age, group and the interaction between age and group were fixed effects (scaled age was introduced in the model) and the random intercept and slope were also considered in the model. In the white matter lesions volume analysis four outliers were excluded (three typical older adults and a superager). White matter volume and white matter lesions volume were adjusted by total intracranial volume (TIV). *P*, *p*-value; SD, standard deviation; SE, standard error.

|  | White matter volume  (TIV-adjusted) | | White matter lesions volume (TIV-adjusted) | | Fazekas score | |
| --- | --- | --- | --- | --- | --- | --- |
|  | **β (SE)** | ***P*** | **β (SE)** | ***P*** | **β (SE)** | ***P*** |
| **Group** | -4.62 (6.11) | 0.45 | 0.38 (0.47) | 0.42 | -0.07 (0.14) | 0.63 |
| **Age (scaled)** | -18.58 (1.13) | < 0.0001 | 0.69 (0.14) | < 0.0001 | 0.10 (0.04) | 0.007 |
| **Group x Age** | 0.40 (1.73) | 0.81 | 0.33 (0.21) | 0.11 | 0.01 (0.05) | 0.80 |
| **Superager slope**, cm^3^/one SD of scaled age | -18.58 (1.13) | - | 0.69 (0.14) | - | 0.11 (0.03) | - |
| **Typical older adult slope**, cm^3^/one SD of scaled age | -18.18 (1.30) | - | 1.01 (0.15) | - | 0.10 (0.04) | - |

**Figure 3-1. ROI-based longitudinal trajectories of white matter fractional anisotropy (FA).** Longitudinal group differences were studied in 18 white matter tracts or regions of interest (ROIs) from the JHU-ICBM atlas. Individual trajectories are plotted together with the group average predicted trajectory. Shaded areas indicate 95% confidence interval. A linear mixed effects model was used to predict average FA in each of the ROIs with group, scaled age and the interaction between the two as fixed factors, the random intercept and slope were included in the model. Age was scaled in the statistical model, but raw values are shown for illustration purposes. All ROIs show a significant (*P* < 0.05) interaction between age and group (indicated with an asterisk (*)). ATR, anterior thalamic radiation; CST, corticospinal tract; Hippo. cing., hippocampal cingulum; IFO, inferior fronto-occipital fasciculus; ILF, inferior longitudinal fasciculus; L, left; R, right; SLF, superior longitudinal fasciculus.

**Figure 3-2. ROI-based longitudinal analysis of white matter fractional anisotropy (FA).** Longitudinal group differences were studied in 18 white matter tracts or regions of interest (ROIs) from the JHU-ICBM atlas with a linear mixed effects model to predict average FA in each of the ROIs with group, scaled age and the interaction between the two as fixed factors, the random intercept and slope were included in the model. ATR, anterior thalamic radiation; β, coefficients; Corr. *P*, false-discovery-rate corrected *p*-value; CST, corticospinal tract; Hippo. cing., hippocampal cingulum; IFO, inferior fronto-occipital fasciculus; ILF, inferior longitudinal fasciculus; L, left; *P*, *p*-value; R, right; SE, standard error; SLF, superior longitudinal fasciculus.

|  | | Group | | | | | | Age (scaled) | | | | | | Group x age (scaled) | | | | | | Slope superagers | | Slope typical older adults |
| --- | --- | --- | --- | --- | --- | --- | --- | --- | --- | --- | --- | --- | --- | --- | --- | --- | --- | --- | --- | --- | --- | --- |
|  |  | β (SE) | | *P* | | Corr. *P* | | β (SE) | | *P* | | Corr. *P* | | β (SE) | | *P* | | Corr. *P* | | estimate (SE) | | estimate (SE) |
| **ATR L.** | -0.013 (0.004) | | 0.002 | | 0.01 | | -0.004 (0.001) | | 3.3x10^-5^ | | 5.9x10^-5^ | | -0.005 (0.001) | | 6.5x10^-5^ | | 0.0003 | | -0.0037 (0.0009) | | -0.0091 (0.001) | |
| **ATR R.** | -0.012 (0.004) | | 0.006 | | 0.03 | | -0.006 (0.001) | | 8.3x10^-16^ | | 3.7x10^-15^ | | -0.003 (0.001) | | 0.004 | | 0.004 | | -0.0060 (0.0007) | | -0.0092 (0.0008) | |
| **Cingulum L.** | -0.007 (0.005) | | 0.17 | | 0.21 | | -0.007 (0.001) | | 3.2x10^-12^ | | 9.6x10^-12^ | | -0.006 (0.002) | | 0.0001 | | 0.0003 | | -0.0070 (0.001) | | -0.0129 (0.0012) | |
| **Cingulum R.** | -0.008 (0.005) | | 0.11 | | 0.19 | | -0.008 (0.001) | | <2.2x10^-16^ | | 1.3x10^-15^ | | -0.005 (0.001) | | 0.0005 | | 0.001 | | -0.0082 (0.0009) | | -0.0129 (0.001) | |
| **CST L.** | -0.001 (0.004) | | 0.75 | | 0.79 | | 3.5x10^-4^ (7.3x10^-4^) | | 0.64 | | 0.67 | | -0.003 (0.001) | | 0.002 | | 0.003 | | 0.0003 (0.0007) | | -0.0031 (0.0009) | |
| **CST R.** | -6.4x10^-4^ (4.0x10^-3^) | | 0.87 | | 0.87 | | -1.2x10^-5^ (7.9x10^-4^) | | 0.99 | | 0.99 | | -4.1x10^-3^ (1.2x10^-3^) | | 0.0007 | | 0.001 | | -0.0001 (0.0008) | | -0.0041 (0.0009) | |
| **Forceps major** | -0.012 (0.005) | | 0.02 | | 0.06 | | -0.009 (9.7x10^-4^) | | <2.2x10^-16^ | | 1.3x10^-15^ | | -0.004 (0.001) | | 0.003 | | 0.004 | | -0.0087 (0.001) | | -0.0130 (0.0011) | |
| **Forceps minor** | -0.008 (0.004) | | 0.03 | | 0.07 | | -0.007 (6.2x10^-4^) | | <2.2x10^-16^ | | 1.3x10^-15^ | | -0.003 (9.6x10^-4^) | | 0.001 | | 0.002 | | -0.0066 (0.0006) | | -0.0097 (0.0007) | |
| **IFO L.** | -0.008 (0.004) | | 0.06 | | 0.11 | | -0.003 (0.001) | | 0.0004 | | 0.0007 | | -0.005 (0.001) | | 0.002 | | 0.003 | | -0.0035 (0.001) | | -0.0081 (0.0011) | |
| **IFO R.** | -0.008 (0.004) | | 0.06 | | 0.11 | | -0.006 (7.5x10^-4^) | | 1.2x10^-14^ | | 4.3x10^-14^ | | -0.003 (0.001) | | 0.007 | | 0.008 | | -0.0058 (0.0007) | | -0.0089 (0.0009) | |
| **ILF L.** | -0.012 (0.004) | | 0.002 | | 0.01 | | 0.007 (0.001) | | 3.7x10^-9^ | | 9.5x10^-9^ | | -0.010 (0.002) | | 2.3x10^-8^ | | 4.1x10^-7^ | | 0.0064 (0.0011) | | -0.0030 (0.0013) | |
| **ILF R.** | -0.011 (0.004) | | 0.01 | | 0.04 | | -0.004 (7.4x10^-4^) | | 5.1x10^-9^ | | 1.2x10^-8^ | | -0.002 (0.001) | | 0.05 | | 0.05 | | -0.0044 (0.0007) | | -0.0066 (0.0009) | |
| **SLF L.** | -0.004 (0.004) | | 0.35 | | 0.39 | | -0.002 (0.001) | | 0.03 | | 0.03 | | -0.005 (0.001) | | 9.6x10^-5^ | | 0.0003 | | -0.0020 (0.0009) | | -0.0073 (0.001) | |
| **SLF R.** | -0.004 (0.004) | | 0.29 | | 0.35 | | -0.004 (0.001) | | 0.006 | | 0.008 | | -0.007 (0.002) | | 0.0006 | | 0.002 | | -0.0037 (0.0013) | | -0.0104 (0.0015) | |
| **Uncinate L.** | -0.013 (0.004) | | 0.0008 | | 0.01 | | 0.001 (0.002) | | 0.42 | | 0.47 | | -0.007 (0.002) | | 0.003 | | 0.004 | | 0.0012 (0.0016) | | -0.0058 (0.0018) | |
| **Uncinate R.** | -0.010 (0.004) | | 0.01 | | 0.04 | | 0.003 (9.4x10^-4^) | | 0.002 | | 0.003 | | -0.008 (0.001) | | 1.1x10^-7^ | | 9.9x10^-7^ | | 0.0029 (0.0009) | | -0.0048 (0.0011) | |
| **Hippo. cing. L.** | -0.007 (0.005) | | 0.13 | | 0.19 | | 0.003 (0.001) | | 0.02 | | 0.02 | | -0.008 (0.002) | | 6.4x10^-6^ | | 3.8x10^-5^ | | 0.0028 (0.0012) | | -0.0056 (0.0014) | |
| **Hippo. cing. R.** | -0.008 (0.006) | | 0.14 | | 0.20 | | -0.005 (0.001) | | 2.6x10^-5^ | | 5.2x10^-5^ | | -0.005 (0.002) | | 0.01 | | 0.01 | | -0.0054 (0.0013) | | -0.0103 (0.0015) | |

**Figure 3-3. ROI-based longitudinal trajectories of white matter mean diffusivity (MD).** Longitudinal group differences were studied bilaterally in 18 white matter tracts or regions of interest (ROIs) from the JHU-ICBM atlas. Individual trajectories are plotted together with the group average predicted trajectory. Shaded areas indicate 95% confidence interval. A linear mixed effects model was used to predict average MD in each of the ROIs with group, scaled age and the interaction between the two as fixed factors, the random intercept and slope were included in the model. Age was scaled in the statistical model, but raw values are shown for illustration purposes. ROIs where the interaction between age and group is significant (*P* < 0.05) are indicated with an asterisk (*), otherwise as non-significant (n.s.). ATR, anterior thalamic radiation; CST, corticospinal tract; Hippo. cing., hippocampal cingulum; IFO, inferior fronto-occipital fasciculus; ILF, inferior longitudinal fasciculus; L, left; R, right; SLF, superior longitudinal fasciculus.

**Figure 3-4. ROI-based longitudinal analysis of white matter mean diffusivity (MD).** Longitudinal group differences were studied in 18 white matter tracts or regions of interest (ROIs) from the JHU-ICBM atlas with a linear mixed effects model to predict average MD in each of the ROIs with group, scaled age and the interaction between the two as fixed factors, the random intercept and slope were included in the model. ATR, anterior thalamic radiation; β, coefficients; Corr. *P*, false-discovery-rate corrected *p*-value; CST, corticospinal tract; Hippo. cing., hippocampal cingulum; IFO, inferior fronto-occipital fasciculus; ILF, inferior longitudinal fasciculus; L, left; *P*, *p*-value; R, right; SE, standard error; SLF, superior longitudinal fasciculus.

|  | Group | | | Age (scaled) | | | | Group x age (scaled) | | | | Slope superagers | | Slope typical older adults | |
| --- | --- | --- | --- | --- | --- | --- | --- | --- | --- | --- | --- | --- | --- | --- | --- |
|  | β (SE) | *P* | Corr. *P* | | β (SE) | *P* | Corr. *P* | | β (SE) | *P* | Corr. *P* | | estimate (SE) | | estimate (SE) |
| **ATR L.** | 6.7x10^-5^ (2.5x10^-5^) | 0.007 | 0.02 | | 3.1x10^-5^ (4.9x10^-6^) | 5.1x10^-10^ | 2.9x10^-9^ | | 2.3x10^-5^ (7.4x10^-6^) | 0.002 | 0.001 | | 3.1x10^-10^ (4.9x10^-11^) | | 5.3x10^-10^ (5.5x10^-11^) |
| **ATR R.** | 5.7x10^-5^ (3.0x10^-5^) | 0.06 | 0.07 | | 3.8x10^-5^ (6.1x10^-6^) | 4.3x10^-10^ | 2.9x10^-9^ | | 2.6x10^-5^ (9.2x10^-6^) | 0.004 | 0.01 | | 3.9x10^-10^ (6.2x10^-11^) | | 6.5x10^-10^ (6.8x10^-11^) |
| **Cingulum L.** | 1.9x10^-5^ (5.1x10^-6^) | 0.0002 | 0.004 | | 8.2x10^-8^ (1.4x10^-6^) | 0.95 | 0.95 | | 6.7x10^-6^ (2.2x10^-6^) | 0.002 | 0.001 | | 2.0x10^-12^ (1.4x10^-11^) | | 6.7x10^-11^ (1.6x10^-11^) |
| **Cingulum R.** | 2.1x10^-5^ (7.0x10^-6^) | 0.003 | 0.02 | | 3.9x10^-6^ (1.8x10^-6^) | 0.03 | 0.04 | | 8.7x10^-6^ (2.8x10^-6^) | 0.002 | 0.01 | | 4.0x10^-11^ (1.8x10^-11^) | | 1.3x10^-10^ (2.1x10^-11^) |
| **CST L.** | 9.0x10^-6^ (5.5x10^-6^) | 0.10 | 0.11 | | 4.3x10^-6^ (1.5x10^-6^) | 0.005 | 0.007 | | -1.0x10^-6^ (2.4x10^-6^) | 0.66 | 0.70 | | 4.3x10^-11^ (1.5x10^-11^) | | 3.2x10^-11^ (1.8x10^-11^) |
| **CST R.** | 6.0x10^-6^ (4.8x10^-6^) | 0.21 | 0.22 | | 1.0x10^-5^ (1.8x10^-6^) | 1.3x10^-8^ | 3.9x10^-8^ | | -3.5x10^-6^ (2.8x10^-6^) | 0.21 | 0.25 | | 1.0x10^-10^ (1.8x10^-11^) | | 6.4x10^-11^ (2.1x10^-11^) |
| **Forceps major** | 4.1x10^-5^ (1.6x10^-5^) | 0.01 | 0.03 | | 6.7x10^-6^ (2.1x10^-6^) | 0.002 | 0.003 | | 8.0x10^-6^ (3.2x10^-6^) | 0.01 | 0.02 | | 6.7x10^-11^ (2.1x10^-11^) | | 1.5x10^-10^ (2.5x10^-11^) |
| **Forceps minor** | 2.1x10^-5^ (8.9x10^-6^) | 0.02 | 0.03 | | 1.1x10^-5^ (1.7x10^-6^) | 8.0x10^-10^ | 2.9x10^-9^ | | 4.1x10^-6^ (2.7x10^-6^) | 0.12 | 0.16 | | 1.1x10^-10^ (1.7x10^-11^) | | 1.5x10^-10^ (2.0x10^-11^) |
| **IFO L.** | 1.9x10^-5^ (7.9x10^-6^) | 0.02 | 0.03 | | 7.6x10^-6^ (2.9x10^-6^) | 0.009 | 0.01 | | 9.0x10^-6^ (4.4x10^-6^) | 0.04 | 0.06 | | 7.7x10^-11^ (2.9x10^-11^) | | 1.7x10^-10^ (3.2x10^-11^) |
| **IFO R.** | 1.7x10^-5^ (8.2x10^-6^) | 0.04 | 0.05 | | 9.6x10^-6^ (1.6x10^-6^) | 7.0x10^-10^ | 2.9x10^-9^ | | 9.1x10^-6^ (2.4x10^-6^) | 0.0001 | 0.002 | | 9.7x10^-11^ (1.5x10^-11^) | | 1.9x10^-10^ (1.8x10^-11^) |
| **ILF L.** | 2.2x10^-5^ (6.7x10^-6^) | 0.0009 | 0.008 | | 2.2x10^-6^ (1.5x10^-6^) | 0.13 | 0.14 | | 4.8x10^-6^ (2.3x10^-6^) | 0.03 | 0.05 | | 2.3x10^-11^ (1.5x10^-11^) | | 6.9x10^-11^ (1.7x10^-11^) |
| **ILF R.** | 1.5x10^-5^ (6.2x10^-6^) | 0.02 | 0.03 | | 3.2x10^-6^ (1.6x10^-6^) | 0.05 | 0.06 | | 7.0x10^-6^ (2.5x10^-6^) | 0.005 | 0.01 | | 3.2x10^-11^ (1.6x10^-11^) | | 1.0x10^-10^ (1.9x10^-11^) |
| **SLF L.** | 2.1x10^-5^ (8.7x10^-6^) | 0.02 | 0.03 | | 1.0e-05 (1.9e-06) | 6.8x10^-8^ | 1.8x10^-7^ | | 5.6x10^-6^ (2.9x10^-6^) | 0.05 | 0.07 | | 1.0x10^-10^ (1.9x10^-11^) | | 1.6x10^-11^ (2.2x10^-11^) |
| **SLF R.** | 2.1x10^-5^ (8.5x10^-6^) | 0.02 | 0.03 | | 7.0x10^-6^ (3.0x10^-6^) | 0.02 | 0.03 | | 1.2x10^-5^ (4.5x10^-6^) | 0.006 | 0.01 | | 7.2x10^-11^ (3.0x10^-11^) | | 1.9x10^-10^ (3.3x10^-11^) |
| **Uncinate L.** | 3.7x10^-5^ (1.5x10^-5^) | 0.02 | 0.03 | | 3.2x10^-5^ (6.2x10^-6^) | 2.1x10^-7^ | 4.7x10^-7^ | | 1.0x10^-5^ (9.3x10^-6^) | 0.26 | 0.29 | | 3.2x10^-10^ (6.2x10^-11^) | | 4.2x10^-10^ (6.9x10^-11^) |
| **Uncinate R.** | -1.1x10^-5^ (1.8x10^-5^) | 0.55 | 0.55 | | 4.3x10^-5^ (4.0x10^-6^) | <2.0x10^-16^ | 3.6x10^-15^ | | 9.6x10^-7^ (6.1x10^-6^) | 0.88 | 0.88 | | 4.3x10^-10^ (4.0x10^-11^) | | 4.4x10^-10^ (4.6x10^-11^) |
| **Hippo. cing. L.** | 2.1x10^-5^ (7.5x10^-6^) | 0.005 | 0.02 | | 7.6x10^-6^ (2.3x10^-6^) | 0.001 | 0.002 | | 9.8x10^-6^ (3.6x10^-6^) | 0.007 | 0.01 | | 7.9x10^-11^ (2.3x10^-11^) | | 1.7x10^-10^ (2.7x10^-11^) |
| **Hippo. cing. R.** | 2.7x10^-5^ (1.2x10^-5^) | 0.03 | 0.04 | | 1.3x10^-5^ (3.1x10^-6^) | 5.0x10^-5^ | 1.0x10^-4^ | | 1.3x10^-5^ (4.8x10^-6^) | 0.005 | 0.01 | | 1.3x10^-10^ (3.1x10^-11^) | | 2.6x10^-10^ (3.6x10^-11^) |
